# Effects of chronodisruption and alcohol consumption on gene expression in reward-related brain areas in female rats

**DOI:** 10.1101/2024.06.19.599697

**Authors:** C. Meyer, K. Schoettner, S. Amir

**Affiliations:** Center for Studies in Behavioral Neurobiology Department of Psychology Concordia University Montreal, QC, H4B 1R6 Canada

**Keywords:** Clock genes, alcohol, females, gene expression, neuroinflammation

## Abstract

Circadian dysfunction caused by exposure to aberrant light-dark conditions is associated with abnormal alcohol consumption in humans and animal models. Changes in drinking behavior have been linked to alterations in clock gene expression in reward-related brain areas, which could be attributed to either the effect of chronodisruption or alcohol. To date, however, the combinatory effect of circadian disruption and alcohol on brain functions is less understood. Moreover, despite known sex differences in alcohol drinking behavior, most research has been carried out on male subjects only, and therefore implications for females remain unclear.

To address this gap, adult female rats housed under an 11h/11h light-dark cycle (LD22) or standard light conditions (LD24, 12h/ 12h light-dark) were given access to an intermittent alcohol drinking protocol (IA20%) to assess the impact on gene expression in brain areas implicated in alcohol consumption and reward: the prefrontal cortex (PFC), nucleus accumbens (NAc), and dorsal striatum (DS). mRNA expression of core clock genes (*Bmal1*, *Clock*, *Per2*), sex hormone receptors (*ERβ*, *PR*), glutamate receptors (*mGluR5*, *GluN2B*), a calcium-activated channel (*Kcnn2*), and an inflammatory cytokine (*TNF-α*) were measured at two-time points relative to the locomotor activity cycle.

Housing under LD22 did not affect alcohol intake but significantly disrupted circadian activity rhythms and reduced locomotion. Significant changes in the expression of *Bmal1*, *ERβ*, and *TNF-α* were primarily related to the aberrant light conditions, whereas changes in *Per2* and *PR* expression were associated with the effect of alcohol. Collectively, these results indicate that disruption of circadian rhythms and/or intermittent alcohol exposure have distinct effects on gene expression in the female brain, which may have implications for the regulation of alcohol drinking, addiction, and, ultimately, brain health.

## INTRODUCTION

Alcohol use disorder is a global public health concern, contributing to millions of premature deaths each year (World Health Organization, 2018). Various factors influence alcohol consumption, including circadian rhythm disruption. Research has shown that working night shifts or rotating schedules can increase the risk of alcohol abuse (Richter et al., 2021), while genetic variations in clock genes are linked to alcohol dependence and increased alcohol intake (Kovanen et al., 2010). Long-term alcohol consumption also interferes with the circadian system, which can further promote alcohol intake (Spanagel, Rosenwasser, et al., 2005). To understand the interplay between circadian rhythm disruption and alcohol consumption, and their impact on health, it is crucial to address their sole and combined effects on the organism.

The circadian system comprises a network of biological clocks located in almost every cell of the body generating daily rhythms in metabolism, physiology, and behavior based on the interaction of so-called clock genes and their proteins. Transcriptional activators Brain and muscle arnt-like 1 (BMAL1) and Circadian locomotor output cycles kaput (CLOCK), and repressors Period (PERs) and Cryptochrome (CRYs) form the core of a transcription-translation feedback loop (TTFL). In brief, heterodimers of BMAL1/CLOCK induce the transcription of *Per* and *Cry* genes, which dimerize once translated into proteins and translocate back to the nucleus where they inhibit the transcriptional activity of BMAL1/CLOCK. Degradation of PER and CRY allows BMAL1/CLOCK to activate transcription again, thus starting a new cycle that repeats every 24 hours. Chronodisruption caused by working in rotating shifts, amongst others, disorganizes the core loop and thus the expression of so-called clock-controlled genes, which mediate the clock output and regulate daily oscillations of physiological and behavioral programs (Bozek et al., 2009).

In the past, both environmental and genetic manipulations have been applied in animal models to study the effects of circadian disruption on alcohol-drinking behavior. Interestingly, exposure to experimental light-dark (LD) conditions aimed to induce misalignment of circadian rhythms, such as chronic jetlag paradigms, revealed inconsistent results. While some studies reported an increase in alcohol intake in male Sprague-Dawley and female HAD-1 rats (Clark et al., 2007; Gauvin et al., 1997), others observed no changes or reduced alcohol intake in female and male Lewis, Fisher, and Wistar rats (Clark et al., 2007; Meyer et al., 2022; Rosenwasser et al., 2010). The use of various animal models and experimental protocols may explain the degree of variability in the observed outcomes, specifically with regard to the level of circadian disruption. Although studies targeting components of the molecular clockwork revealed clear associations between the expression of single clock genes and alcohol consumption, responses are not uniform across different clock genes. While global knockouts of *Per2* and *Clock* have been shown to increase alcohol consumption in mice (Ozburn et al., 2013; Spanagel, Pendyala, et al., 2005), deficiencies in *Rev-erbα* and *Cry1/2* have been associated with reduced alcohol preference and consumption (Al-Sabagh et al., 2022; Hühne et al., 2022). As such, it is conceivable that the results of studies using environmentally induced chronodisruption vary depending on the level of disorganized clock gene expression.

The circadian system is sexually dimorphic (Yan & Silver, 2016) and sex differences in alcohol consumption in animals with disrupted circadian rhythms have been reported (Rizk et al., 2022; Rosenwasser et al., 2014). Moreover, there is emerging evidence that circadian clock genes in specific brain regions mediate sex differences in alcohol-drinking behavior (de Zavalia et al., 2022). However, because the impact of chronodisruption on alcohol consumption has been studied in males primarily, implications for females remain unclear.

Also, unknown to date is the impact of alcohol consumption on circadian clock function in brain regions regulating alcohol-drinking behavior. Alcohol has been found to affect the circadian system at the molecular and behavioral level (Brager et al., 2010; Chen et al., 2004; Logan et al., 2010; Rosenwasser et al., 2005), which may exacerbate the effects of chronodisruption, thus promoting abnormal alcohol consumption or the development of alcohol use disorder depending on sex.

To address this, we tested the distinct and combined effects of chronodisruption and alcohol consumption on gene expression in specific brain regions that have been identified to regulate drinking behavior in rodents. While female rats did not consume more alcohol when chronodisrupted, our results indicate that the combination of exposure to aberrant light conditions and alcohol consumption modified the expression of clock genes and genes associated with proinflammatory responses in the rats’ brains, whereas the isolated effects of chronodisruption or alcohol were less prominent. Our results contribute to the understanding of the molecular consequences of circadian disruption and/or alcohol use in female rats and may help in the development of targeted interventions and recommendations to treat alcohol abuse in females.

## RESULTS

### Locomotor Activity and Circadian Rhythm Parameter

We first assessed the locomotor activity of rats kept under LD24, LD22, and in combination with the IA20% paradigm. Visual inspection of the actograms showed that exposure to aberrant light conditions affected locomotor activity rhythms, whereas alcohol did not (Figure 1).

**Figure 1:**
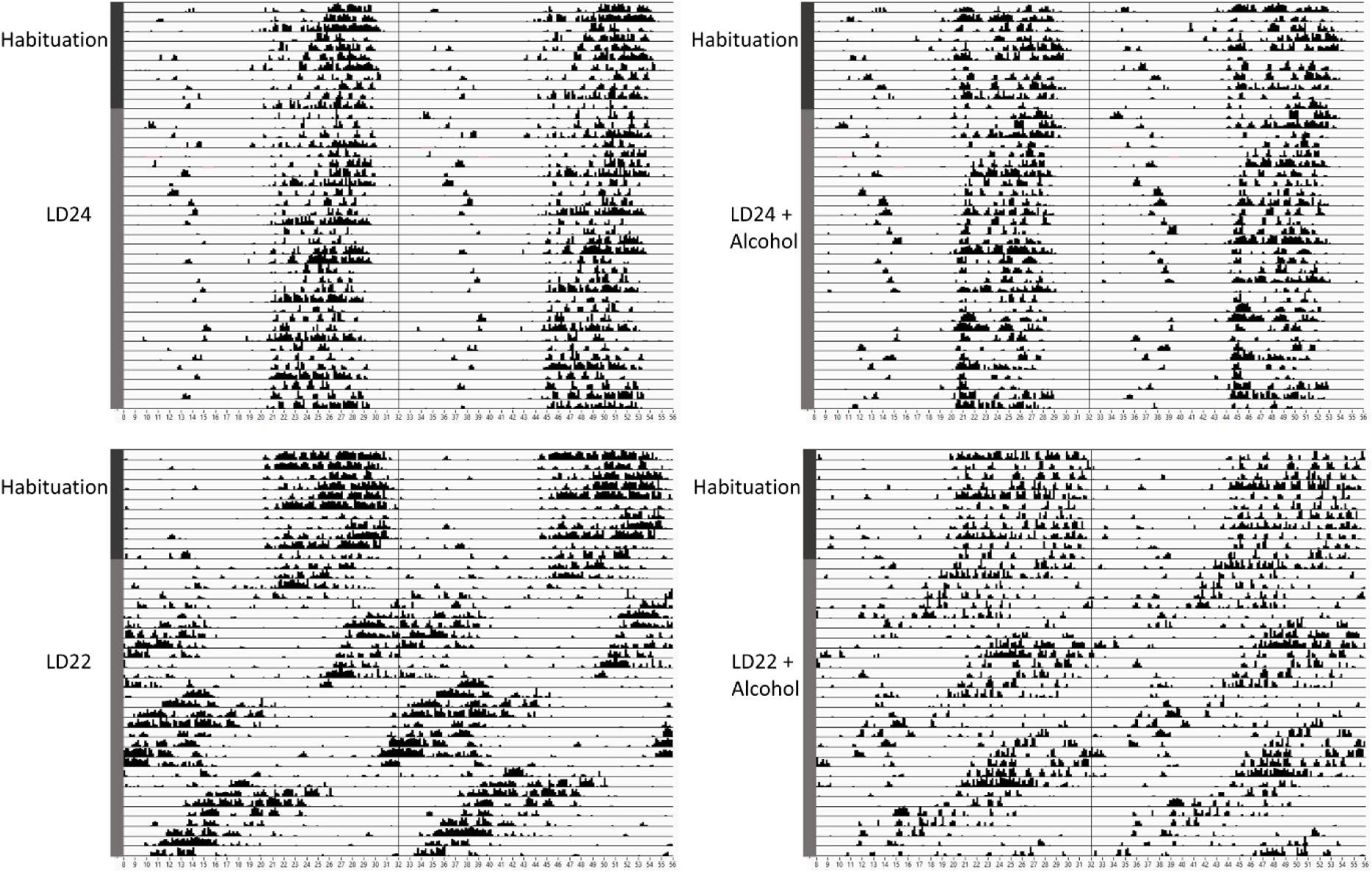
Representative double-plotted actograms of locomotor activity rhythms of female rats kept under various light conditions (standard LD12:12 h or aberrant LD11:11 h cycle) and intermittent access to alcohol solution (IA20%) over 40 days. The time of day is double-plotted along the x-axis, and successive days are arranged from top to bottom along the y-axis.

As expected, the circadian parameters Interdaily Stability (IS) and Relative Amplitude (RA) differed between the LD groups (Figure 2; One-way ANOVA, IS: F(3,20)= 11.78, p= 0.0001; RA: F(3,20)= 10.13 p= 0.0003). However, Intradaily Variability (IV) did not vary between LD24 and LD22 (IV: F(3,20)= 0.6710, p= 0.5798). Alcohol consumption did not affect the assessed circadian parameters.

**Figure 2:**
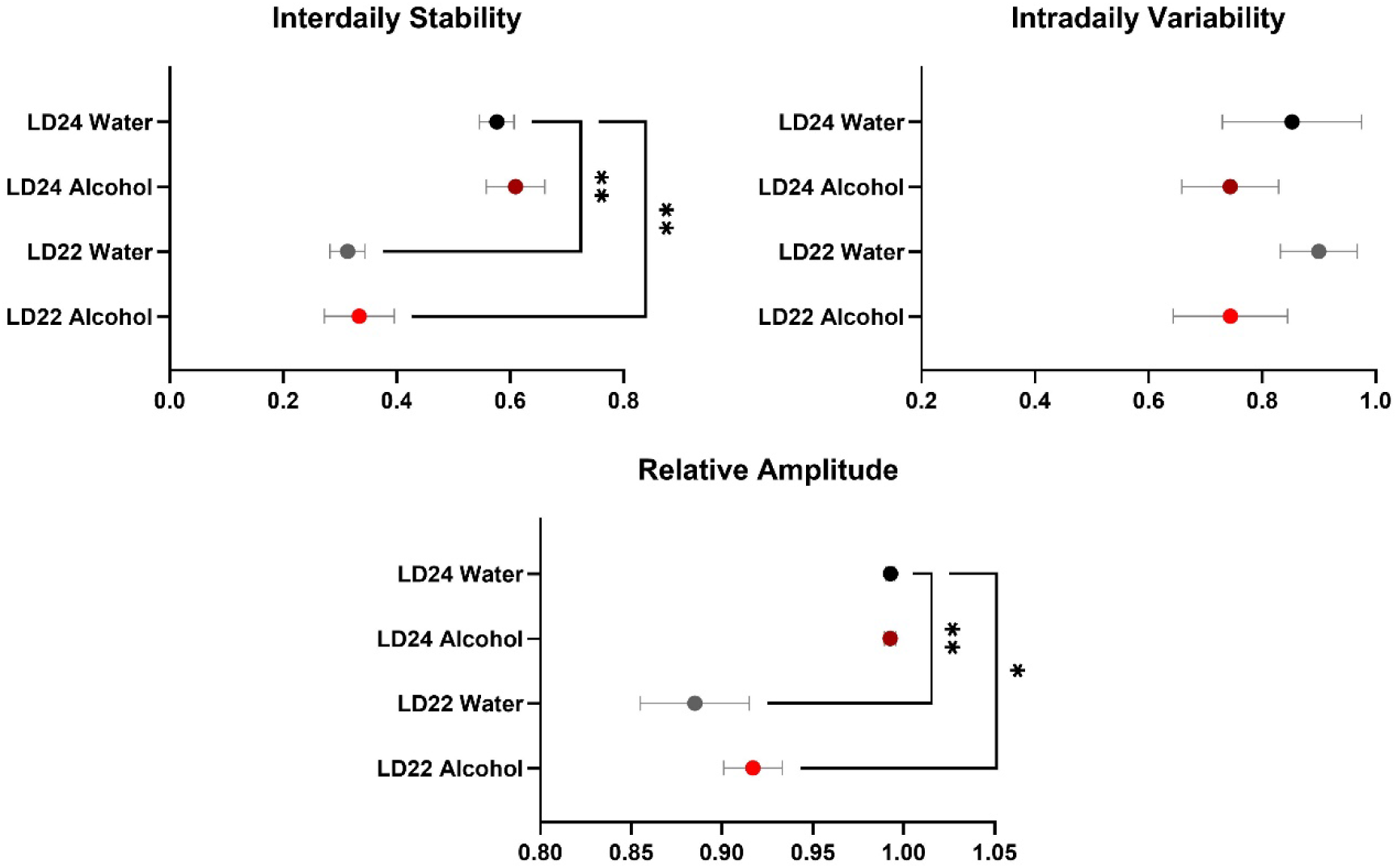
Circadian rhythm parameters during the last 10 days of the experiment. The Interdaily Stability (IS) and Relative Amplitude (RA) of the circadian rhythms were significantly affected by the LD condition, while Intradaily Variability (IV) was unchanged. Data are presented as mean ± SEM, with n= 6 animals per LD and fluid condition. One-way ANOVA followed by Tukey’s multiple comparisons test: * p < 0.05, ** p < 0.01.

Further analysis revealed significant suppression of locomotor activity under the LD22 condition, but no effect of alcohol (Figure 3; Two-way ANOVA: Fluid Condition p= 0.4676; LD Condition p= 0.0041, Interaction p= 0.7811).

**Figure 3:**
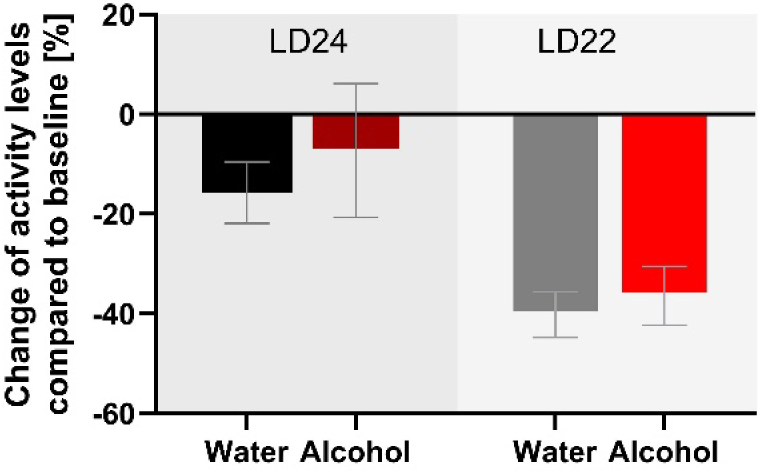
Exposed to aberrant light conditions suppress locomotion while alcohol does not affect activity levels. The sum of activity counts recorded in 10-minute intervals over eight days is shown. Data are presented as mean ± SEM, with n= 6 animals per LD and fluid condition.

### Alcohol Drinking Behavior

All rats, regardless of LD condition, significantly increased their alcohol intake and preference over the course of the intermittent alcohol exposure sessions (Figure 4, Alcohol consumption: Session F(3.975, 39.75)= 6.383 p= 0.0005; Alcohol preference: Session F(4.898, 48.98)= 4.566 p= 0.0018). However, we did not observe any differences in alcohol intake or preference between the LD24 and LD22 groups (Two-way ANOVA; LD F(1, 10)= 1.160, p= 0.3068) and alcohol preference (Two-way ANOVA: LD F(1, 10)= 0.3119, p= 0.5888).

**Figure 4:**
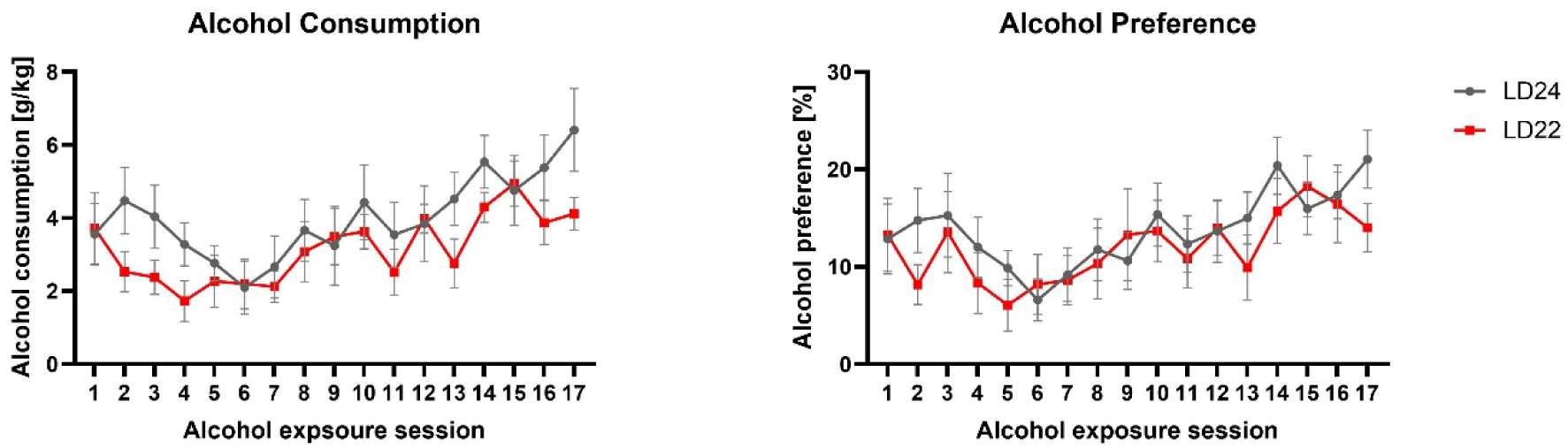
Alcohol consumption and preference were unaffected in female rats exposed to aberrant light conditions. Rats in both the LD24 and LD22 groups showed similar drinking patterns that changed significantly as the experiment progressed. Data are presented as mean ± SEM, with n= 6 animals per LD condition.

### Gene Expression

Using real-time polymerase chain reaction (PCR), we examined the gene expression changes of three key clock genes (*Bmal1*, *Per2*, and *Clock*) in the prefrontal cortex (PFC), nucleus accumbens (NAc), and dorsal striatum (DS) of rats exposed to different LD and fluid conditions (Figure 5, Table 1).

**Figure 5:**
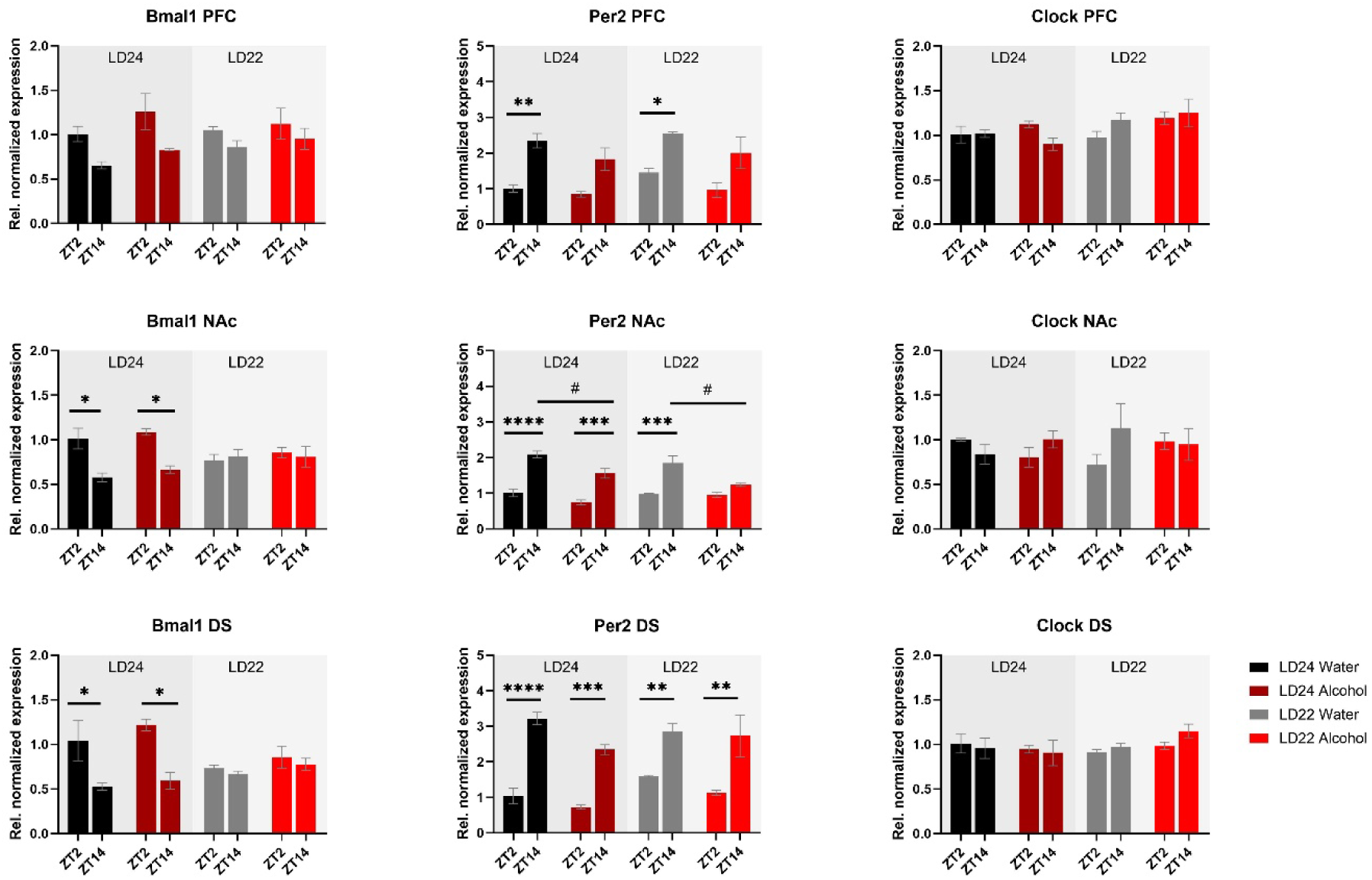
mRNA expression of clock genes *Bmal1*, *Per2*, and *Clock* in the PFC, NAc, and DS at ZT2 and ZT14. The expression of *Bmal1* and *Per2,* but not *Clock* was affected by the light or fluid condition differently (see Results section and Table 1 for details). Data are presented as mean ± SEM, with n= 3 animals per LD condition and fluid condition. Three-way ANOVA with Šídák’s multiple comparisons test: **** p < 0.0001, *** p < 0.001, ** p < 0.01, * p < 0. 05.

**Table 1:** Statistical analysis of gene expression.

| Target | Brain Region | Three-way ANOVA |  |  | Interactions |
| --- | --- | --- | --- | --- | --- |
|  |  | LD Condition | Fluid Condition | ZT |  |
| *Bmal1* | PFC | n.s. (p= 0.4655) | n.s. (p= 0.0825) | **p= 0.0024** | ZT x LD Condition: n.s. (p= 0.2100)<br>ZT x Fluid Condition: n.s. (p= 0.8646)<br>LD Condition x Fluid Condition: n.s. (p= 0.4204)<br>ZT x LD Condition x Fluid Condition: n.s. (p= 0.7512) |
|  | NAc | n.s. (p= 0.6497) | n.s. (p= 0.2625) | **p= 0.0011** | ZT x LD Condition **p= 0.0011**<br>ZT x Fluid Condition: n.s. (p= 0.7389)<br>LD Condition x Fluid Condition: n.s. (p= 0.7260)<br>ZT x LD Condition x Fluid Condition: n.s. (p= 0.6195) |
|  | DS | n.s. (p= 0.2562) | n.s. (p= 0.1312) | **p= 0.0005** | ZT x LD Condition **p= 0.0041**<br>ZT x Fluid Condition: n.s. (p= 0.6825)<br>LD Condition x Fluid Condition: n.s. (p= 0.9883)<br>ZT x LD Condition x Fluid Condition: n.s. (p= 0.7479) |
| *Per2* | PFC | n.s. (p= 0.1609) | **p= 0.0167** | **p< 0.0001** | ZT x LD Condition: n.s. (p= 0.7798)<br>ZT x Fluid Condition: n.s. (p= 0.5426)<br>LD Condition x Fluid Condition: n.s. (p= 0.5904)<br>ZT x LD Condition x Fluid Condition: n.s. (p= 0.6400) |
|  | NAc | n.s. (p= 0.2296) | **p= 0.0003** | **p< 0.0001** | ZT x LD Condition **p= 0.0284**<br>ZT x Fluid Condition **p= 0.0162**<br>LD Condition x Fluid Condition: n.s. (p= 0.6452)<br>ZT x LD Condition x Fluid Condition: n.s. (p= 0.3095) |
|  | DS | n.s. (p= 0.1049) | **p= 0.0059** | **p< 0.0001** | ZT x LD Condition: n.s. (p= 0.1211)<br>ZT x Fluid Condition: n.s. (p= 0.6872)<br>LD Condition x Fluid Condition: n.s. (p= 0.3003)<br>ZT x LD Condition x Fluid Condition: n.s. (p= 0.1332) |
| *Clock* | PFC | **p= 0.0418** | n.s. (p= 0.2656) | n.s. (p= 0.8571) | ZT x LD Condition: n.s. (p= 0.0774)<br>ZT x Fluid Condition: n.s. (p= 0.1474)<br>LD Condition x Fluid Condition: n.s. (p= 0.2531)<br>ZT x LD Condition x Fluid Condition: n.s. (p= 0.7353) |
|  | NAc | n.s. (p= 0.7350) | n.s. (p= 0.8923) | n.s. (p= 0.3318) | ZT x LD Condition: n.s. (p= 0.4200)<br>ZT x Fluid Condition: n.s. (p= 0.8517)<br>LD Condition x Fluid Condition: n.s. (p= 0.7967)<br>ZT x LD Condition x Fluid Condition: n.s. (p= 0.0632) |
|  | DS | n.s. (p= 0.4188) | n.s. (p= 0.5801) | n.s. (p= 0.5889) | ZT x LD Condition: n.s. (p= 0.1986)<br>ZT x Fluid Condition: n.s. (p= 0.6346)<br>LD Condition x Fluid Condition: n.s. (p= 0.1472)<br>ZT x LD Condition x Fluid Condition: n.s. (p= 0.7148) |
| *ERβ* | PFC | **p= 0.0415** | n.s. (p= 0.5835) | **p= 0.0437** | ZT x LD Condition: n.s. (p= 0.4458)<br>ZT x Fluid Condition: n.s. (p= 0.7292)<br>LD Condition x Fluid Condition: n.s. (p= 0.1893)<br>ZT x LD Condition x Fluid Condition: n.s. (p= 0.6341) |
|  |  | LD Condition | Fluid Condition | ZT |  |
|  | NAC | n.s. (p= 0.1352) | n.s. (p= 0.2899) | n.s. (p= 0.2338) | ZT x LD Condition: n.s. (p= 0.8456)<br>ZT x Fluid Condition: n.s. (p= 0.3247)<br>LD Condition x Fluid Condition: n.s. (p= 0.5881)<br>ZT x LD Condition x Fluid Condition: n.s. (p= 0.2671) |
|  | DS | **p= 0.0030** | n.s. (p= 0.4058) | n.s. (p= 0.1459) | ZT x LD Condition: n.s. (p= 0.5813)<br>ZT x Fluid Condition: n.s. (p= 0.1776)<br>LD Condition x Fluid Condition: n.s. (p= 0.3637)<br>ZT x LD Condition x Fluid Condition: n.s. (p= 0.6530) |
| PR | PFC | n.s. (p= 0.6542) | **p= 0.0483** | n.s. (p= 0.6584) | ZT x LD Condition: n.s. (p= 0.7862)<br>ZT x Fluid Condition: n.s. (p= 0.1362)<br>LD Condition x Fluid Condition: n.s. (p= 0.3492)<br>ZT x LD Condition x Fluid Condition: n.s. (p= 0.6347) |
|  | NAC | n.s. (p= 0.0798) | n.s. (p= 0.5399) | n.s. (p= 0.2874) | ZT x LD Condition: n.s. (p= 0.5353)<br>ZT x Fluid Condition **p= 0.0459**<br>LD Condition x Fluid Condition: n.s. (p= 0.9295)<br>ZT x LD Condition x Fluid Condition: n.s. (p= 0.6730) |
|  | DS | n.s. (p= 0.3889) | n.s. (p= 0.5351) | n.s. (p= 0.1766) | ZT x LD Condition: n.s. (p= 0.9317)<br>ZT x Fluid Condition: n.s. (p= 0.8058)<br>LD Condition x Fluid Condition: n.s. (p= 0.4593)<br>ZT x LD Condition x Fluid Condition: n.s. (p= 0.6474) |
| mGluR5 | PFC | n.s. (p= 0.4126) | n.s. (p= 0.9470) | n.s. (p= 0.1737) | ZT x LD Condition: n.s. (p= 0.6113)<br>ZT x Fluid Condition: n.s. (p= 0.4981)<br>LD Condition x Fluid Condition: n.s. (p= 0.6221)<br>ZT x LD Condition x Fluid Condition: n.s. (p= 0.2912) |
|  | NAC | n.s. (p= 0.5843) | n.s. (p= 0.2183) | n.s. (p= 0.0766) | ZT x LD Condition: n.s. (p= 0.2917)<br>ZT x Fluid Condition **p= 0.0039**<br>LD Condition x Fluid Condition **p= 0.0390**<br>ZT x LD Condition x Fluid Condition: n.s. (p= 0.1359) |
|  | DS | n.s. (p= 0.9071) | n.s. (p= 0.3266) | n.s. (p= 0.3422) | ZT x LD Condition: n.s. (p= 0.4925)<br>ZT x Fluid Condition: n.s. (p= 0.6652)<br>LD Condition x Fluid Condition **p= 0.0387**<br>ZT x LD Condition x Fluid Condition: n.s. (p= 0.7295) |
| GluN2B | PFC | n.s. (p= 0.4958) | n.s. (p= 0.1562) | n.s. (p= 0.0610) | ZT x LD Condition: n.s. (p= 0.8064)<br>ZT x Fluid Condition: n.s. (p= 0.3543)<br>LD Condition x Fluid Condition: n.s. (p= 0.0713)<br>ZT x LD Condition x Fluid Condition: n.s. (p= 0.8853) |
|  | NAC | n.s. (p= 0.5085) | n.s. (p= 0.2876) | n.s. (p= 0.4104) | ZT x LD Condition: n.s. (p= 0.4687)<br>ZT x Fluid Condition: n.s. (p= 0.2654)<br>LD Condition x Fluid Condition: n.s. (p= 0.7785)<br>ZT x LD Condition x Fluid Condition: n.s. (p= 0.5495) |
|  | DS | n.s. (p= 0.5381) | n.s. (p= 0.4703) | n.s. (p= 0.3017) | ZT x LD Condition: n.s. (p= 0.8562)<br>ZT x Fluid Condition: n.s. (p= 0.2523)<br>LD Condition x Fluid Condition: n.s. (p= 0.0695)<br>ZT x LD Condition x Fluid Condition: n.s. (p= 0.9955) |
| Kcnn2 | PFC | **p= 0.0471** | n.s. (p= 0.7844) | n.s. (p= 0.3668) | ZT x LD Condition: n.s. (p= 0.7811)<br>ZT x Fluid Condition: n.s. (p= 0.2633)<br>LD Condition x Fluid Condition: n.s. (p= 0.7233)<br>ZT x LD Condition x Fluid Condition: n.s. (p= 0.6586) |
|  | NAC | n.s. (p= 0.2198) | n.s. (p= 0.3944) | **p= 0.0352** | ZT x LD Condition: n.s. (p= 0.3744)<br>ZT x Fluid Condition: n.s. (p= 0.8438)<br>LD Condition x Fluid Condition: n.s. (p= 0.9572)<br>ZT x LD Condition x Fluid Condition: n.s. (p= 0.8113) |
|  | DS | n.s. (p= 0.1695) | n.s. (p= 0.3149) | n.s. (p= 0.8818) | ZT x LD Condition: n.s. (p= 0.7157)<br>ZT x Fluid Condition: n.s. (p= 0.7579) |
|  |  | LD Condition | Fluid Condition | ZT |  |
|  |  |  |  |  | LD Condition x Fluid Condition: n.s. (p= 0.7579)<br>ZT x LD Condition x Fluid Condition: n.s. (p= 0.7485) |
| *TNF-α* | PFC | **p= 0.0040** | n.s. (p= 0.5850) | n.s. (p= 0.5784) | ZT x LD Condition: n.s. (p= 0.3050)<br>ZT x Fluid Condition: n.s. (p= 0.6730)<br>LD Condition x Fluid Condition: n.s. (p= 0.7818)<br>ZT x LD Condition x Fluid Condition: n.s. (p= 0.0970) |
|  | NAc | **p= 0.0158** | n.s. (p= 0.9826) | **p= 0.0421** | ZT x LD Condition: n.s. (p= 0.4758)<br>ZT x Fluid Condition: n.s. (p= 0.2660)<br>LD Condition x Fluid Condition: n.s. (p= 0.6879)<br>ZT x LD Condition x Fluid Condition: n.s. (p= 0.6142) |
|  | DS | **p= 0.0054** | n.s. (p= 0.2668) | **p= 0.0359** | ZT x LD Condition: n.s. (p= 0.6412)<br>ZT x Fluid Condition: n.s. (p= 0.0643)<br>LD Condition x Fluid Condition: n.s. (p= 0.2793)<br>ZT x LD Condition x Fluid Condition: n.s. (p= 0.7507) |

Expression patterns of clock genes *Bmal1* and *Per2* in the PFC, NAc, and DS were significantly affected by time of day, whereas the effects of photoperiods (LD24 vs LD22) and alcohol vary depending on the gene (Figure 5, Table 1). A notable reduction in the amplitude of *Bmal1* expression measured at ZT2 and ZT14 was observed in the striatum of animals kept under LD22 cycles, while alcohol consumption did not change levels of *Bmal1* expression in either brain region. Interestingly, the photoperiod did not affect *Per2* expression levels, demonstrated by noticeable differences in *Per2* levels at ZT2 and ZT14 that were present under both LD22 and LD24 photoperiods (Figure 5 center panel, Table 1). *Per2* expression in the PFC, DS, and most notably in the NAc, however, was affected by the fluid condition animals were exposed to (Figure 5 center panel, Table 1). In contrast, no effects of photoperiod, time of day, or alcohol exposure on Clock gene expression were found, except for the PFC where *Clock* appears marginally upregulated under the LD22 light cycle (Figure 5 right panel, Table 1).

mRNA expression of the hormone receptors *ERβ* and *PR* was further examined (Figure 6). While expression of both receptors was not markedly affected by the experimental conditions, three-way ANOVA revealed changes in *ERβ* levels in the PFC depending on the time of day, while photoperiod affected *ERβ* in the PFC and DS (Table 1). *PR* mRNA expression in the PFC was marginally but significantly affected by the fluid condition (Table 1) while remaining unchanged in the dorsal and ventral striatum irrespective of time of day, photoperiod, or drinking paradigm.

**Figure 6:**
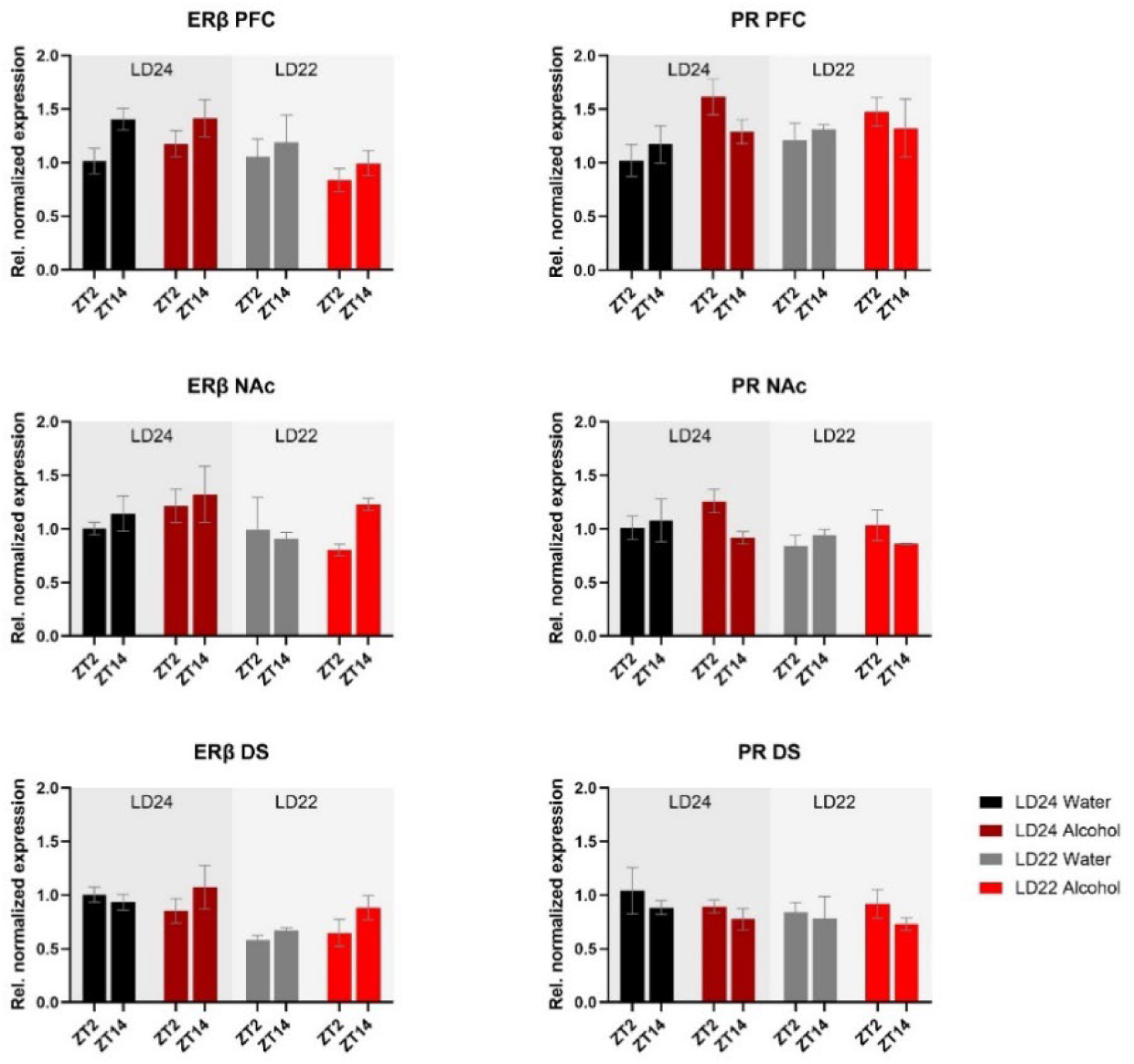
Expression of Estrogen Receptor β (*ERβ*) and Progesterone Receptor (*PR*) in the PFC, NAc, and DS at ZT2 and ZT14. The mRNA expression of *ERβ* and *PR* was only mildly affected by the LD condition or fluid condition (see Results section and Table 1 for details). Results are presented as mean ± SEM, with n= 3 animals per LD and fluid condition.

Furthermore, we investigated genes related to glutamate signaling, including the glutamate receptors *mGluR5* and *GluN2B*, and the small conductance calcium-activated potassium channel protein 2 (*Kcnn2*) (Figure 7). No main effects of time of day, photoperiod, or drinking paradigm were observed in either brain region on *mGluR5* and *GluN2B*, while *Kcnn2* expression in the PFC and NAc was marginally affected by the light cycle or time of day, respectively (Table 1).

**Figure 7:**
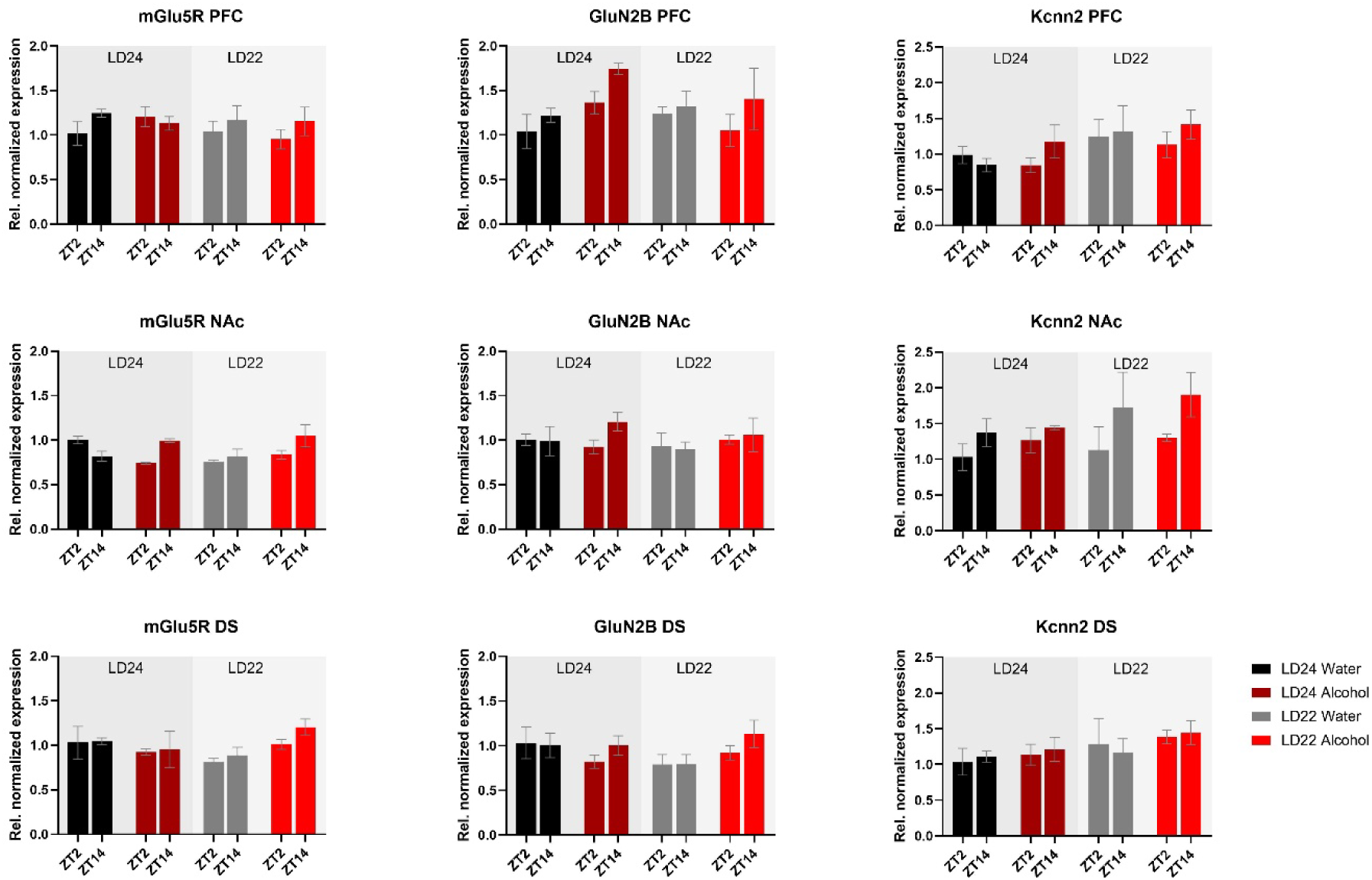
Expression of glutamate receptors (*mGluR5*, *GluN2B*) and the Calcium-Activated Potassium Channel (*Kcnn2*) in the PFC, NAc, and DS at ZT2 and ZT14. The mRNA expression of the glutamate receptor *mGluR5* was affected by the interaction of LD and fluid conditions, while no changes were observed in the *GluN2B* receptor. mRNA levels of *Kcnn2* were slightly increased under the LD22 condition in the PFC and NAc (see Results section and Table 1 for details). Results are presented as mean ± SEM, with n= 3 animals per LD condition and fluid condition.

The expression of inflammatory marker *TNF-α* was also examined (Figure 8). Statistical analysis revealed significant main effects of photoperiod on expression patterns in all three brain regions (Table 1), suggesting that aberrant light conditions led to an increase in *TNF-α* expression. While time of day had a significant effect on striatal *TNF-α* expression as well, alcohol consumption appears to exacerbate the difference although no significant interaction was found (Table 1).

**Figure 8:**
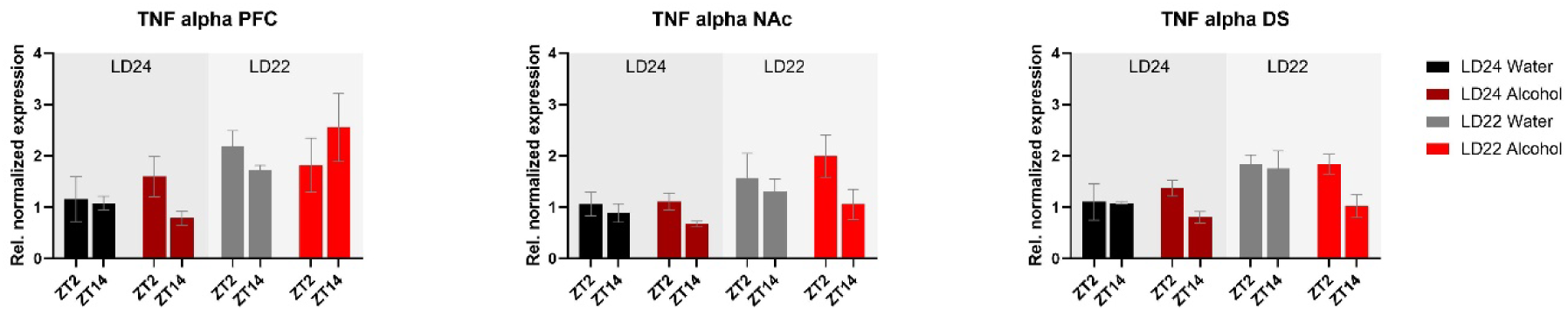
Expression of the inflammatory marker *TNF-α* in the PFC, NAc, and DS at ZT2 and ZT14. The mRNA expression of *TNF-α* was affected by both the LD22 condition and time of day. Results are presented as mean ± SEM, with n= 3 animals per LD condition and fluid condition.

## DISCUSSION

Disruption of circadian rhythms caused by exposure to abnormal light-dark (LD) conditions is considered a risk factor for alcohol abuse (Barko et al., 2019). However, the neurobiological mechanisms underlying this relationship, particularly in females, are not well understood. In this study, we examined the impact of circadian disruption and intermittent alcohol consumption on the expression of genes in brain areas involved in alcohol-drinking behavior.

Consistent with previous findings, housing female rats under LD22 conditions resulted in the desynchronization of locomotor activity rhythms, but it had no significant effect on alcohol consumption or preference (Meyer et al., 2022). At the molecular level, expression patterns of *Bmal1* and *TNF-α* in the striatum were mostly affected by the aberrant light conditions, whereas alcohol significantly affected the expression of *Per2* in the PFC, NAc, and DS only.

Overall, the results indicate that in females, aberrant LD conditions and/or alcohol consumption have only mild effects on the expression of clock and cell-signaling genes within reward-related brain areas. This may explain why alcohol-drinking behavior remains similar under both standard and altered LD conditions in female rats. Based on these findings, it can be concluded that, in female rats, chronodisruption is not a main factor leading to abnormal alcohol-drinking behavior.

### Circadian Rhythmicity and Locomotor Activity

Changes in circadian rhythm parameters have been associated with neurodegenerative diseases (Anderson et al., 2009; Hatfield et al., 2004). Consistent with previous findings in male rats (De La Iglesia et al., 2004), our study confirmed that the LD22 schedule exhibits chronodisruptive characteristics in females. However, despite a significant decline in daily activity and consolidated rest-activity patterns, rhythm fragmentation remained unchanged irrespective of photoperiod and fluid conditions. The reduced rhythmicity resulted from impaired day-to-day coordination, evidenced by decreased interdaily stability (IS) and dampened robustness of activity rhythms (RA), rather than a fragmentation within each day (unchanged intradaily variability, IV) (Banks et al., 2022).

We further assessed the levels of wheel running activity, as previous research has demonstrated that changes in circadian photoperiod and alcohol exposure can influence locomotor activity (Benstaali et al., 2002; Perreau-Lenz & Spanagel, 2015). Our findings revealed a significant decline in activity under LD22 conditions, while alcohol access did not affect the levels of wheel running. Prior studies have reported conflicting effects of non-standard LD conditions on locomotion, with both increases and decreases observed in male rodents (Bartoszewicz et al., 2009; Delorme et al., 2022; Okuliarova et al., 2016). In female mice, both forward and inverted (shift work-like) photoperiod shifts were found to decrease locomotion (Banks et al., 2022). Notably, the reported changes in locomotor activity were not associated with any significant alterations in alcohol-drinking behavior under experimental LD conditions (Gamsby & Gulick, 2015; Rosenwasser et al., 2010). Hence, the employed experimental LD condition in this study may have a more pronounced impact on locomotor behavior compared to the alcohol exposure paradigm, and the altered locomotor behavior does not appear to affect alcohol intake.

### Alcohol Drinking Behavior

The intermittent alcohol exposure (IAE20%) paradigm used in this study involves daily cycles of alcohol drinking and abstinence, which mimics the repetitive pattern of excessive intake, abstinence, and relapse observed in individuals with alcohol abuse and dependence (Crabbe et al., 2011; Koob & Volkow, 2016), which induces pharmacologically relevant blood alcohol concentrations in rats (Carnicella et al., 2014).

Consistent with earlier studies using the IA20% paradigm in male rats (Mill et al., 2013; Simms et al., 2008), the female rats in our study also exhibited increased alcohol consumption over time. However, we did not observe an effect of an aberrant light cycle on alcohol intake behavior, thus confirming the results of previous studies (Meyer et al., 2022; Rizk et al., 2022).

### Gene Expression

Non-standard light conditions have been shown to disrupt rhythms of clock gene expression in various tissues (Oishi et al., 2015; Szántóová et al., 2011) and affect the diurnal regulation of reward-related processes (Depoy et al., 2017). Furthermore, alcohol exposure has been reported to alter molecular and behavioral rhythms (Guo et al., 2016) and to affect the expression of genes regulating reward-related processes in the brain (Logan et al., 2014). Through gene expression analysis, we aimed to investigate the interplay of environmental circadian disruption and alcohol consumption on genes involved in circadian function and cell signaling in brain areas mediating reward processes.

Gene association studies in humans have linked the core clock genes *Bmal1*, *Per2*, and *Clock* to abnormal alcohol consumption (Dong et al., 2011; Kovanen et al., 2010), and animal studies have supported these findings. Mutations of *Per1*, *Per2*, and *Clock* in mice have been shown to enhance alcohol consumption, likely through their impact on brain reward processes (Gamsby et al., 2013; Ozburn et al., 2013; Rizk et al., 2022; Spanagel, Pendyala, et al., 2005), whereas *Cry1/2-*deficient mice exhibited decreased alcohol preference (Hühne et al., 2022). Notably, a targeted knockout of *Bmal1* in medium spiny neurons (MSNs) of the striatum decreased alcohol consumption in female mice (de Zavalia et al., 2021), while *Bmal1* ablation in the NAc resulted in higher alcohol consumption (Herrera et al., 2023).

Moreover, the selective deletion of *Per2* in either the entire striatum or the NAc did not affect alcohol consumption in female mice (de Zavalia et al., 2021; Herrera et al., 2023). The results suggest that clock genes may have inhibitory or stimulatory effects on alcohol consumption in females, depending on the specific clock gene and brain region, highlighting the intricate regulation of alcohol consumption by these genes.

Our data indicated differential effects of aberrant light conditions and alcohol consumption on the mRNA expression of the clock genes *Clock*, *Bmal1*, and *Per2* in female rats. Interestingly, *Clock* expression remained unaffected by photoperiod and alcohol exposure when compared to control animals, which may explain the similar drinking patterns in both groups. Previous findings by Ozburn et al. (2013) showed that mice bearing a *Clock* knockout exhibited significantly increased alcohol intake, particularly in females, while others reported that female C*lock*-deficient and WT female mice consumed similar amounts of alcohol, unlike males (Rizk et al., 2022). Notably, these drinking phenotypes remained unchanged when animals were kept under aberrant LD schedules (Rizk et al., 2022). Despite the contradictory outcomes in studies on *Clock* knockout mice that may be attributed to variations in drinking paradigms or the use of different mouse strains, our results confirm the view that robust expression of *Clock*, particularly in the NAc (Ozburn et al., 2013), may confer protection from abnormal drinking, among other factors.

The expression of *Bmal1* appeared to be dysregulated in the striatum and prefrontal cortex in animals exposed to aberrant light conditions but their drinking remained unchanged compared to control animals. This suggests that, in females, the sole abundance of *Bmal1* in the striatum rather than its oscillation plays a role in the regulation of alcohol consumption as demonstrated by experiments in mice with ablation of *Bmal1* from neurons of the entire striatum or just the NAc (de Zavalia et al., 2021; Herrera et al., 2023).

It could also be suggested that certain clock-controlled processes may remain largely unaffected by the dysregulation of *Bmal1* expression, as indicated by the levels of *Per2* mRNA in the mPFC, NAc, and DS in females kept under LD22 photoperiods, potentially mitigating the effect of chronodisruption on alcohol consumption. Although the persistence of differences in *Per2* mRNA levels at ZT2 and ZT14 under LD22 conditions were unexpected, they align with trends observed in other studies. For example, male and female mice with a striatum-specific deletion of *Bmal1* show only mild changes in *Per2* expression in the dorsal striatum (Schoettner et al., 2022). Additionally, male mice exposed to aberrant LD conditions display disrupted *Per2* expression in the SCN, while robust expression profiles of *Per2* were observed in the striatum (Ikeno & Yan, 2016). These findings suggest that other factors, such as dopamine signaling, may influence the expression of clock genes in the mesolimbic system, as demonstrated by Hood et al. (2010). Despite recent advances, further studies are needed to better understand the interaction of cell signaling pathways, such as the dopaminergic circuit, the circadian clock, or individual clock genes in the reward system, and how they influence drug-related behaviors.

Previous studies have associated *Per2* expression with alcohol consumption in rodents, but recent findings in mice suggest that these effects are sex-dependent. For example, a global or striatum-specific knockout of *Per2* increased alcohol consumption in male mice (de Zavalia et al., 2021; Herrera et al., 2023; Spanagel, Pendyala, et al., 2005). In contrast, female mice with striatum-specific *Per2* knockout display no change in alcohol consumption, indicating that other sex-specific factors must be considered in the regulation of alcohol-drinking behavior in females.

Sex hormones could potentially be the link between sex-dependent alcohol consumption and clock gene expression in reward-regulating brain areas. Previous studies have shown that CLOCK-BMAL1 complexes bind to the E-box element in the promoter region of estrogen receptor β (*ERβ*), thereby influencing its expression and subsequently altering cell signaling pathways, such as glutamatergic signaling in the striatum of female rats (Grove-Strawser et al., 2010), which may indirectly affect alcohol consumption (Pandey et al., 2004). In our study, we found that the levels of *ERβ* mRNA remained unchanged irrespective of LD conditions, levels of *Bmal1* expression, or access to alcohol. To determine, whether estrogen signaling is affected by exposure to aberrant LD conditions and/ or alcohol exposure, future investigations should include a functional analysis and explore the levels of fluctuating sex hormones.

It is also widely recognized that changes in glutamatergic circuits have an impact on reward-related behaviors, including alcohol consumption, and vice versa (Chandrasekar, 2013; Eisenhardt et al., 2015; Gass & Olive, 2008). Numerous studies have established a connection between the metabotropic glutamate receptor subtype-5 (*mGluR5*) and the regulation of various behavioral pathologies associated with alcohol misuse. Previous research has demonstrated that alcohol consumption modulates *mGluR5* in a brain-region-specific manner (Cozzoli et al., 2009; Simonyi et al., 2004) and that *mGluR5* receptor manipulations affect alcohol intake (Cozzoli et al., 2014; Goodwani et al., 2017).

Furthermore, the inotropic NMDA receptor serves as a key target for the actions of alcohol within the central nervous system, and behavioral and cellular investigations have pointed to the importance of the *GluN2B* subunit in mediating the effects of alcohol (Allgaier, 2002; Nagy et al., 2005). NMDA receptors are implicated in synaptic plasticity and are involved in various aspects of drug and alcohol addiction (Abrahao et al., 2013; Kroener et al., 2012). However, the effects of alcohol on NMDA receptors are inconsistent, as acute alcohol exposure has been shown to inhibit, while chronic exposure promotes the function of the *GluN2B* subunit (Nagy et al., 2003; Sheela Rani & Ticku, 2006; Wang et al., 2007).

Additionally, small-conductance calcium-activated potassium channels (SKs) modulate NMDA receptor-dependent synaptic plasticity (Faber et al., 2005; Lin et al., 2008) and facilitate learning and memory acquisition (Hammond et al., 2006; Sun et al., 2020). Previous research has shown that chronic alcohol exposure in mice led to a reduction in the expression of *Kcnn2* (KCa2.2) and a significant increase in the expression of *GluN2B* (Mulholland et al., 2011).

Overall, our study did not reveal substantial changes in mRNA expression of the selected neurotransmitter receptors mentioned above following chronic alcohol consumption and/ or chronodisruption. While this aligns with the observed lack of changes in alcohol-drinking behavior, further work needs to evaluate their functions using electrophysiological approaches to draw final conclusions, as gene expression analysis may not directly relate to receptor density and function.

In addition to their influence on cell signaling, a growing body of evidence supports the pivotal role of the circadian clock genes in the regulation of inflammatory processes (Carter et al., 2015), which may contribute to alterations in alcohol consumption. Studies have shown that the deletion of *Bmal1* and *Rev-erbα* leads to widespread glial activation, inflammation, and oxidative stress in the brain (Griffin et al., 2019; Musiek et al., 2013). Likewise, chronic alcohol intake has been reported to induce neuroinflammation characterized by increased expression of proinflammatory cytokines, such as tumor necrosis factor-alpha (*TNF-α*) (Blednov et al., 2005, 2011; Cooper et al., 2020; Kelley & Dantzer, 2011). Activation of *TNF-*α has been associated with impaired neuronal functioning and increased risk of alcohol abuse and relapse (Obad et al., 2018). Our findings demonstrate that aberrant light cycles, but not chronic alcohol consumption, amplify *TNF-α* mRNA levels in all examined brain regions. However, *TNF-α* is a nonspecific inflammatory marker, and future studies should investigate the complex relationship between circadian disruption, clock genes, and neuroinflammatory processes in greater detail.

Both, this study, and previous work in mice indicate that female rodents may exhibit greater resilience to aberrant light conditions concerning alcohol consumption. However, a confounding factor in these findings may be the use of running wheels. Exercise is known to influence alcohol-drinking behavior (Buhr et al., 2021; Gallego et al., 2015; Werme et al., 2002). It may be interesting for future studies to compare the effect of aberrant light conditions on alcohol intake and gene expression in the presence and absence of running wheels.

## CONCLUSION

The results of this study show that environmentally induced circadian rhythm disruptions did not affect alcohol consumption in female rats, despite dysregulation of *Bmal1* expression in several brain regions associated with alcohol-drinking behavior. It is possible that other clock genes, such as *Per2*, maintain rhythmicity in brain areas important for reward processing, or that the expression of genes like *Clock* remains unchanged, contributing to compensatory mechanisms to counteract the loss of *Bmal1* rhythmicity.

Our data indicated a more intricate pattern of clock gene expression in the striatum, a brain region that governs alcohol drinking in a sex-dependent manner in rodents, regardless of the presence or absence of specific clock genes. Future studies should focus on investigating specific neural circuits to elucidate the relations between molecular changes and behavioral phenotypes in females.

## MATERIALS AND METHODS

### Ethics Statement

All experimental protocols were approved by the Concordia University Animal Care Committee (certificate number: 30000256).

### Animals and Light-Dark Conditions

Adult female Wistar rats were purchased from Charles River (St. Constant, QC, Canada). Animals were singly housed in clear plexiglass cages (9.5“x8“x16”) equipped with running wheels in light-(standardized illuminance of ∼200–300 lux during the light period, 0 lux during the dark period) and temperature- and humidity-controlled rooms (temperature: 21±1°C, relative humidity: 65±5%). Wheel-running activity of each animal was continuously recorded with the VitalView system (Mini-Mitter, Starr Life Sciences Corp., Oakmont, USA). Food and water were available *ad libitum*. Only animals with a regular estrous cycle were included in the study.

Following a 3-week habituation to the housing and experimental setting, half of the animals were placed under LD22 (11:11 h light-dark, n=12), and the other half remained under standard LD24 (12:12 h light-dark, n=12) conditions. The LD22 light paradigm has been shown to disrupt circadian rhythms and to affect temperature, sleep, corticosterone secretion, and mood (Ben-Hamo et al., 2016; Cambras et al., 2007; De La Iglesia et al., 2004; Wotus et al., 2013).

### Intermittent Alcohol Drinking Paradigm

Female rats kept under LD24 or LD22 were randomly assigned into two groups (n=6) and given access to alcohol (20% v/v, in tap water) every second day for 17 days (IAE20%) as previously described (Wise, 1973). On alcohol days, animals were given access to one bottle of 20% alcohol solution and one bottle of water. After 24 hours, the bottles were replaced with two water bottles that were available for the next 24 hours. This pattern was repeated each Monday, Wednesday, and Friday. Over the weekend, animals had access to water for 48 hours. To control side preferences, the position of the alcohol and water bottles was switched for each alcohol exposure session. Alcohol intake (g/kg body weight/day) and alcohol preference (g of alcohol solution consumed/ g of total fluid intake) were measured and calculated at the end of each daily alcohol session. The remaining animals continued drinking water only throughout the experiment.

### Gene Expression

Gene expression was assessed in tissue punches collected from the prefrontal cortex (PFC), nucleus accumbens (NAc), and dorsal striatum (DS). Animals were euthanized 24h to 72h after the last alcohol exposure at ZT2 (n= 3) and ZT14 (n= 3) (ZT-Zeitgeber time; ZT0 represents the time of light onset). In animals kept under LD22, ZT2 refers to two hours after activity offset and ZT14 to two hours after activity onset. Animals kept under LD24 were euthanized at the stage of proestrus. Animals exposed to LD22 did not show a regular estrous cycle (Meyer et al., 2022). The estrous cycle stage was visually determined by the locomotor activity pattern over four days, as previously described (Wollnik & Turek, 1988).

After euthanasia, brains were flash-frozen and stored at −80°C until further processing. 200μm thick coronal sections were obtained using a cryostat (Microm HM505 E, Microm International GmbH, Walldorf, Germany), and tissue punches (2 mm in diameter) were collected from each hemisphere following the rat brain atlas (Paxinos & Watson, 2006).

Total RNA isolation was conducted following a standard Trizol extraction protocol according to the manufacturer’s instructions (Invitrogen™, Life Technologies Corporation, Carlsbad, CA, United States), and total RNA yield was measured using spectrophotometry (Nanodrop 2000; Thermo Scientific™, Wilmington, DE, United States). Next, the RNA integrity was determined by the “bleach gel” method (Aranda et al., 2012) cDNA was synthesized from 1μg of RNA using the iScriptTMReverse Transcription Supermix following the manufacturer’s instructions (Biorad, Hercules, CA, United States). Besides the standard cDNA samples, a no reverse transcriptase (no-RT) control was prepared.

Realtime PCR was performed with SYBR®Green Supermix SYPR (Biorad, Hercules, CA, United States) and amplification was done using the CFX96TM Real-Time PCR system (Biorad, Hercules, CA, United States). Quantification of target genes was calculated by using the delta-delta Ct (ΔΔCt) method (Pfaffl, 2001) by the CFX Maestro qPCR Software (Biorad, Hercules, CA, United States). The Ct values were normalized to those of β-actin and Gapdh. The primers for the used genes are shown in Table 2.

**Table 2:**
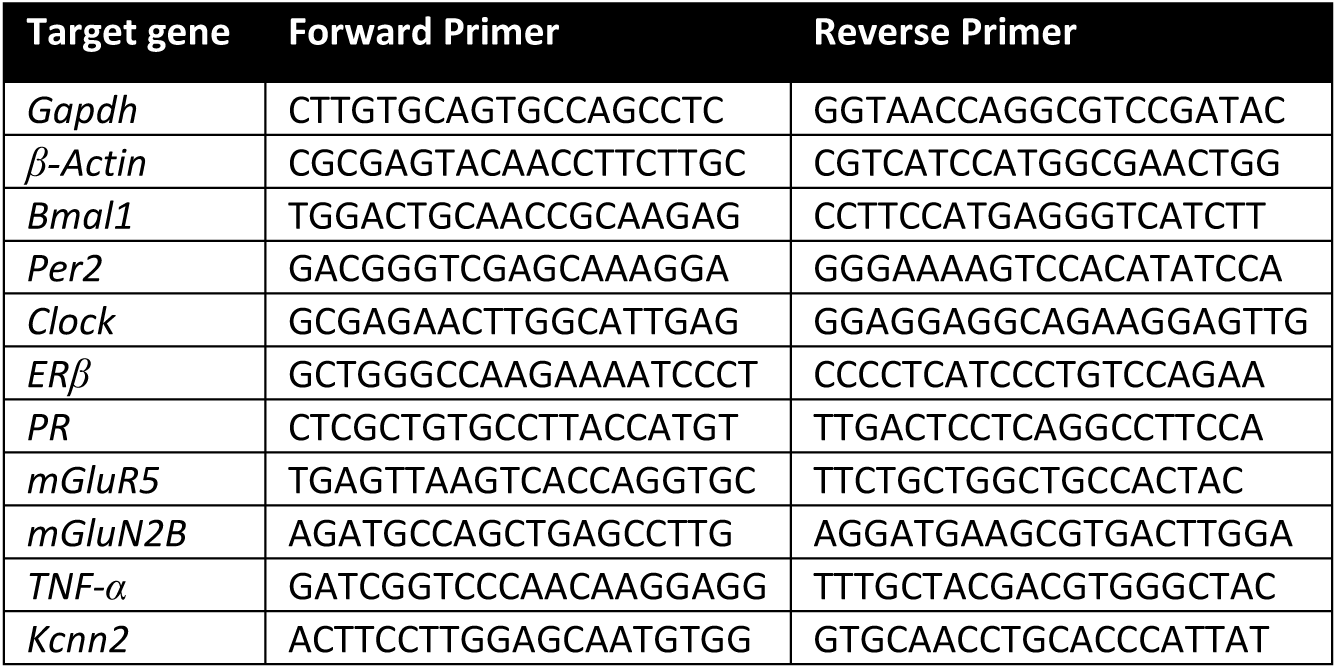
Primer list for quantitative real-time PCR.

### Data Analysis and Statistics

All data from locomotor activity, circadian measurements, and gene expression were analyzed using Prism 9 (GraphPad Software, San Diego, CA, United States). Results are expressed as mean ± standard error of the mean (SEM) and were examined for normality and homogeneity of variance. The significance level was set at p < 0.05.

Circadian rhythmicity was analyzed by continuous registration of locomotor activity using running wheels. Summed wheel revolutions were recorded over 10-minute periods using the VitalView system (Mini-Mitter, Starr Life Sciences Corp., Oakmont, USA). Wheel running activities were visualized as double-plotted actograms using ClockLab 6 (Actimetrics Software, Evanston, United States). The circadian rhythm parameters, Relative Amplitude (RA), Intradaily Variability (IV), and Interdaily Stability (IS), were calculated using ClockLab 6 over the last ten days of the experiment. Differences between the treatment groups were calculated by a one-way ANOVA, followed by Tukey’s multiple comparisons tests.

Locomotor activity counts were determined over eight days during habituation and eight days at the end of the experiment. As locomotor activity varies in female rodents depending on the stage of the estrous cycle, we assessed the activity counts over eight days as a multiple of the four-day estrous cycle. Changes in locomotor activity during treatment compared to baseline were calculated as a percentage.

Alcohol consumption and alcohol preference over water were assessed by a two-way repeated measures ANOVA (LD and alcohol session effect). A three-way ANOVA (LD, fluid condition, and ZT effect) followed by Šídák’s multiple comparisons test was used to analyze the gene expression data.

## AUTHOR CONTRIBUTIONS

The experiments were conceived and designed by CM and SA. Experiments were conducted by CM and assisted by KS. Data were analyzed and interpreted by CM with input from KS and SA. Data visualization was done by CM. The manuscript was written by CM and revised by SA and KS. SA supervised the project. All authors approved the submitted version.

